# Mean fitness is maximized in small populations under stabilizing selection on highly polygenic traits

**DOI:** 10.1101/2025.11.17.688329

**Authors:** Aaron P. Ragsdale

## Abstract

Stabilizing selection commonly acts on complex traits that affect individual fitness. Here, we relate mean fitness under stabilizing selection to population size and trait architecture, using a simple application of theoretical predictions for the distribution of phenotypic values in a single-trait Gaussian stabilizing selection model. We show that mean fitness is maximized by a finite (often small) population size when the total diploid mutation rate *U* across trait-affecting loci is reasonably large. Namely, this occurs when 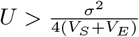, where *σ*^2^ is the variance of the distribution of effect sizes of new mutations, *V*_*S*_ determines the strength of stabilizing selection, and *V*_*E*_ is the environmental variance. We validate these predictions using simulations and briefly discuss their implications for interpreting genetic load and adaptability in small populations.

## Introduction

It is generally understood in conservation and population genetics that a reduced population size poses evolutionary challenges for species survival and individual fitness. In a small population, deleterious variation is allowed to segregate at higher frequencies (Kimura et al., 1963), contributing to genetic load (Haldane, 1937; Crow, 1958), and mildly and moderately deleterious mutations fix more readily, leading to reduced absolute fitness among individuals (Lande, 1994; Lynch et al., 1995a,b). A smaller population size also results in lower levels of standing variation, limiting the response to selection when faced with a novel selective pressure or a changing environment (Lande and Shannon, 1996; Hayward and Sella, 2022).

It has also been established that the mode of selection on individual loci is an important factor in predicted genetic load and standing functional variation. For example, Kimura et al. (1963) showed that mildly deleterious mutations can contribute more to load than strongly deleterious ones, and in some scenarios partially recessive deleterious variation can result in small, finite populations having smaller load than large populations (see also Keller and Waller, 2002). Smaller populations may, in certain scenarios, more effectively purge recessive deleterious variants, as those alleles are more frequently exposed to selection as homozygotes (Kyriazis et al., 2021). Nonetheless, many examples from natural systems, domesticated species, and lab experiments show that reduced long-term *N*_*e*_ corresponds to increased measures of genetic load and reduced overall diversity (e.g., Paland and Schmid, 2003; Willi et al., 2013; Pekkala et al., 2014, and recently reviewed in Robinson et al. (2023)).

In much of this literature, the phenotype of interest is fitness itself. To make model predictions, the selective effects of the many deleterious mutations that reduce fitness are typically combined multiplicatively, and selection acts directionally on individual fitness. However, for many traits that contribute to fitness variation in natural populations, stabilizing selection acts to maintain phenotypic values near some intermediate optimum (Sanjak et al., 2018; Sella and Barton, 2019; Koch et al., 2024; Patel et al., 2025). Instead of directional negative selection on unconditionally deleterious variants, selection at trait-affecting loci resembles underdominance (Robertson, 1956) – stabilizing selection acts to reduce the genetic variance of a trait, which manifests as selection against the minor allele. Selection should be more effective in a large population, keeping allele frequencies relatively rarer and thus lowering the genetic contribution contributed by a given allele. Yet, larger populations have a relatively larger mutational input, so that total genetic variance may increase.

With stabilizing selection on a quantitative trait, how do population size and trait architecture interact to affect mean fitness? Does our intuition still hold, in which small populations have decreased mean fitness? We would expect that stabilizing selection more effectively maintains a population’s mean phenotype near the optimal trait value when the population is large. At the same time, the genetic variance of a trait increases with the population size, and this depends also on the rate and effects of new mutations and the strength of stabilizing selection. Below, we combine these predictions to show that mean fitness is maximized in small populations when the total rate of mutations *U* affecting the trait is sufficiently large, a condition that is likely met for many complex traits of interest.

### Mean fitness with stabilizing selection on one trait

We consider a randomly mating population of diploid individuals with size *N*_*e*_. We model individual fitness following a standard Gaussian stabilizing selection model, with inverse strength of stabilizing selection *V*_*S*_, so that an individual with the optimal phenotype has fitness one. We note that the environmental variance can be absorbed into the fitness function (Lande, 1975, and see Supporting Information (SI) section A), and results are therefore independent of *V*_*E*_.

The fitness of an individual with trait value *z*, relative to the optimal phenotype *z*_0_, is then

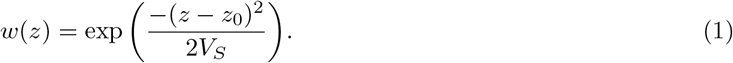

In what follows, we set the optimal phenotype to zero.

Given some distribution *f* (*z*) of phenotypic values among individuals in the population, the mean fitness of the population is found by taking the integral

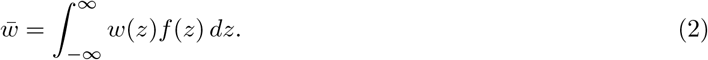

We assume that the distribution of phenotypes is close to (but not necessarily centered on) the optimum. With many mutations contributing to the trait, phenotypes will be approximately normally distributed with variance *V*_*P*_ and mean 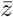: that is, 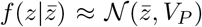. Furthermore, 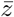 is expected to fluctuate around the optimum, and the displacement from the optimum (*δ*) is normally distributed with mean zero and variance 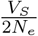 (Lande, 1976; *Simons et al*., *2018): that is*,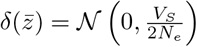.

Then the expected mean fitness of the population, integrating over the distribution of 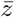, is

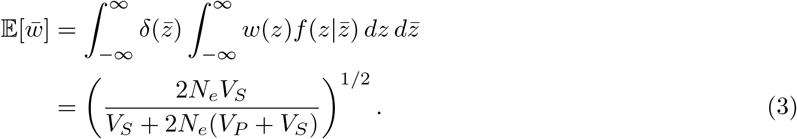

In the limit *N*_*e*_ → ∞, we recover

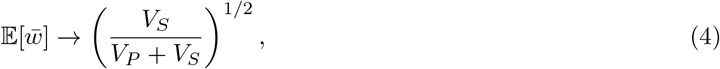

as deviations of the population mean phenotype approach zero with increasing population size.

### The stochastic house of cards model

We assume linkage equilibrium between loci, and we ignore epistasis, dominance and pleiotropy. Under these assumptions, the phenotypic variance is equal to the additive genetic variance, *V*_*G*_. Generally, *V*_*G*_ will depend on the total diploid mutation rate *U* (= 2*µL*, with *µ* the per-base haploid rate, *L* the target size for the trait), the distribution of mutation effect sizes (assumed to be *N* (0, *σ*^2^)), and the strength of selection (determined by *V*_*S*_). Assuming underdominant selection at trait-affecting loci, the mean *V*_*G*_ can be found from the distribution of allele frequencies (e.g., Wright, 1945; Kimura, 1964) and integrating over the distribution of effect sizes, which is handled numerically (SI section B).

A number of approximations for *V*_*G*_ are available from the literature. If effect sizes are large (*σ*^2^ ≫ 1*/N*_*e*_), *V*_*G*_ ≈ 2*UV*_*S*_, a classic result originally due to Latter (1960). In the opposite limit, when effect sizes are weak (*σ*^2^ ≪1*/N*_*e*_), *V*_*G*_ is linear in the population size-scaled mutation rate and variance of effect sizes (Lande, 1975), so *V*_*G*_ ≈ 2*N*_*e*_*Uσ*^2^. The stochastic house of cards (SHOC) model interpolates these two regimes (see Ch. VII in Bürger (2000), Ch. 28 in Walsh and Lynch (2018), and references therein), so that

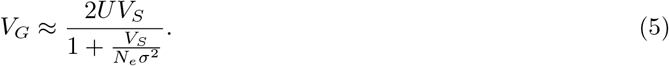

This is a good approximation at steady state across a wide range of parameters, provided *V*_*G*_ ≪*V*_*S*_ and effect sizes are drawn from a normal distribution, though it performs worse for other distributions of effect sizes (Figure S1).

Replacing this expression for *V*_*P*_ in the prediction for 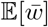 (Eq. 3), we obtain

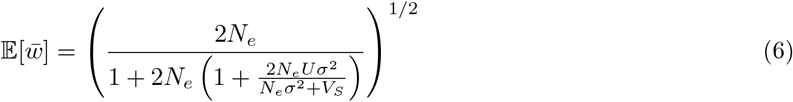

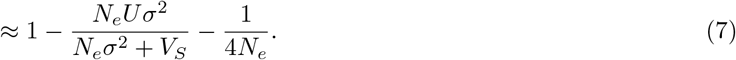

When effects are normally distributed, we find that this prediction (Eq. 6) is accurate across a wide range of model parameters (Fig. 1). For other distributions of effect sizes, while the behavior is qualitatively similar, the SHOC approximation provides a poorer fit to observed *V*_*G*_ and thus average fitness. In general, as *N*_*e*_ becomes large the genetic load approaches *U*, which is the well known expectation for genetic load in a large population, and as *N*_*e*_ becomes small the genetic load is approximately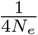, reflecting the drift load.

**Figure 1:**
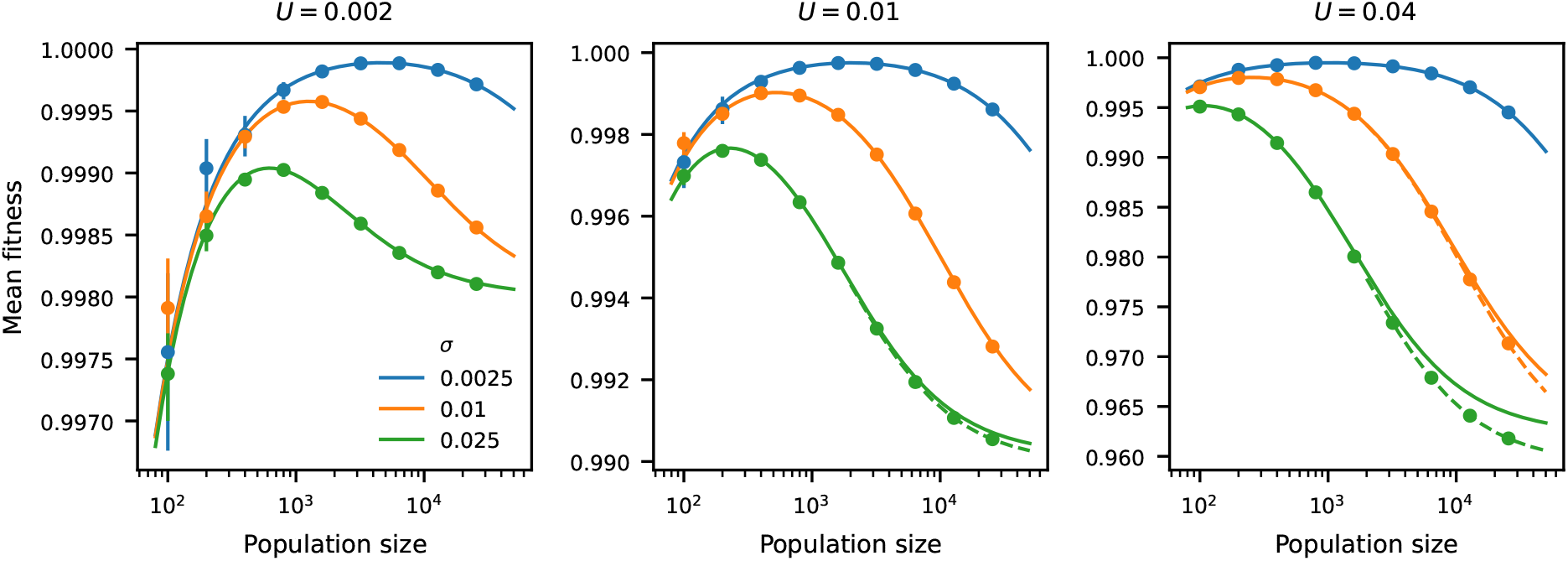
Small population sizes can maximize mean fitness with polygenic traits under stabilizing selection. Predictions for mean fitness within populations (Eq. 6, solid lines) show that mean fitness may decline as the population size increases. For larger mutation rates and effect sizes, *V*_*G*_ may not be small relative to *V*_*S*_, so that Eq. 5 is an underestimate. *V*_*G*_ may instead be computed from the expected distribution of allele frequencies across the distribution of effect sizes and then 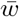 computed from Eq. 3 (dashed lines, SI section B), providing a more accurate prediction in this region of parameter space. Simulations (SI section C) assumed linkage equilibrium across all loci and closely match predictions. Bars display 95% confidence intervals (which are narrower than the marker widths except for small population sizes and mutation rates). Individual-based simulations similarly show close agreement to model predictions (SI section D).

### Condition for increased mean fitness with finite population size

Taking the derivative of 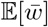 (Eq. 6) with respect to *N*_*e*_ is simple (if a bit tedious), and the numerator is quadratic in *N*_*e*_. We can set it equal to zero and quickly solve for the effective size that maximizes expected mean fitness. This yields a positive solution,

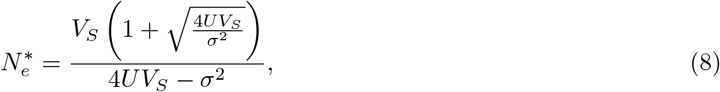

when 4*UV*_*S*_ *> σ*^2^, that is, if the total mutation rate at trait-affecting loci

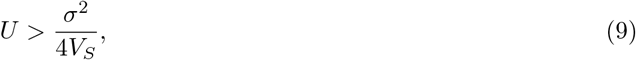

or if the target size is large enough,

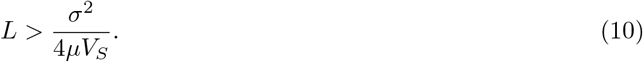

When 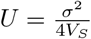, the genetic load is equal to the genetic load in an infinite population (≈*U* ). The difference between the genetic load in a population with size 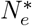 and that in an infinite population grows as *U* becomes larger than 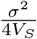, with the difference approaching *U* when 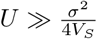 . If the condition (9) is not met, the mean fitness increases monotonically with the population size.

In the presence of uncorrelated environmental noise contributing to phenotypes, we can replace *V*_*S*_ with *V*_*S*_ + *V*_*E*_, so a finite *N*_*e*_ maximizes mean fitness in the population when

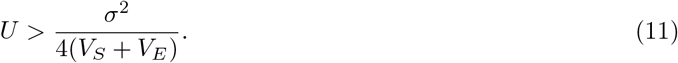

## Conclusions

For traits under stabilizing selection, smaller populations harbor lower genetic variance (Eq. 5), but the average phenotype tends to drift farther from the optimum 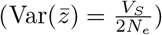. This can be understood through the opposite dependencies of the mutation load and the drift load on *N*_*e*_ (Eq. 6). If the total mutation rate *U* is sufficiently large, then mutation-stabilizing selection balance results in the genetic variance of the trait being high enough in large populations so that the average fitness across all individuals is lower relative to a small or intermediate population sizes.

Are the conditions in which this occurs (Eq. 9) relevant to complex traits in natural populations? In a recent analysis of human GWAS results from 95 traits, Simons et al. (2025) find that *L* and *h*^2^ both vary considerably across traits, but they estimate that *L* is typically of order 10^6^ or larger. Distributions of effect sizes span multiple orders of magnitude, although *σ*^2^ is difficult to estimate for any given trait. We may instead consider the relationship

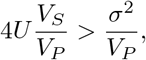

and ask whether we expect it to be satisfied. For a number of human quantitative traits, *V*_*S*_*/V*_*P*_ has been estimated to be ∼ 60 (28–173), implying weak but non-negligible stabilizing selection (Sanjak et al., 2018). If we assume a per-base mutation rate of *µ* ∼ 10^*™*8^ and *L* ≳ 10^6^ (and ignore environmental variance), the quantity on the left should easily exceed 1. And while *σ*^2^ may be unknown, we should expect *σ*^2^*/V*_*P*_ to fall below 1, implying that our condition is likely met.

Such large mutational target sizes for complex traits in humans (with *L*∼ *O* (10^6^) or greater) implies extensive pleiotropy across trait architectures. The model considered here is simple – a single trait under direct stabilizing selection. Even with highly pleiotropic architectures, we note that allele and trait dynamics under pleiotropic stabilizing selection are well-captured by an appropriate single-trait model (Simons et al., 2018; Sella and Barton, 2019), and predictions can be made using the expected distribution of allele frequencies (SI section E). While outside the scope of this note, the effects of pleiotropy on predictions presented here warrant further investigation. Furthermore, direct applications to human trait architectures may be challenging, because neither multivariate trait fitness functions nor the distribution of mutational effects across traits are expected to be isotropic, and their covariances are largely unknown. Here, we have mostly assumed a Gaussian distribution of effects for a single trait, which has been commonly deployed in the literature due to its mathematical convenience. The genetic load may be sensitive to the precise distribution of effect sizes (Figure S1), which is unknown.

Demographic and selection processes are not expected to remain at steady state. In particular, phenotypic optima can change, and a population’s rate of approach to the new optimum is determined by the genetic variance of the trait (Lande, 1994; Hayward and Sella, 2022). A population with reduced genetic variance, as is the case in small populations, may be challenged to readily adapt to a new phenotypic optimum (SI section F, Figure S4), and small population sizes would likely be subject to increased load when the environment fluctuates frequently. Understanding fitness and trait dynamics in a continually changing environment and for non-equilibrium and structured populations is a clear direction for future work (Bertram and Shafiei, 2026; Veller and Muralidhar, 2026).

## Data and code availability

Python scripts and a Mathematica notebook to recreate these results are available at https://github.com/apragsdale/stabilizing_selection_Ne.

## Acknowledgments

I thank K. Thornton, C. Veller and Y. Simons for useful discussions and N. Anderson for helpful comments on an earlier version of this manuscript. This work was supported by the National Institute of General Medical Sciences of the National Institutes of Health (R35GM154962 to APR). Computational resources were generously provided through UW-Madison’s Center for High Throughput Computing.

## References

Jason Bertram and Zahra Shafiei. Strong amplification of quantitative genetic variation under a balance between mutation and fluctuating stabilizing selection. Genetics, 233(1):iyag063, 2026.

Reinhard Bürger. The mathematical theory of selection, recombination, and mutation. John Wiley & Sons, 2000.

James F Crow. Some possibilities for measuring selection intensities in man. Human biology, 30(1):1–13, 1958.

J.B.S. Haldane. The effect of variation on fitness. American Naturalist, 71:337–349, 1937.

Laura Katharine Hayward and Guy Sella. Polygenic adaptation after a sudden change in environment. Elife, 11:e66697, 2022.

Lukas F Keller and Donald M Waller. Inbreeding effects in wild populations. Trends in ecology & evolution, 17(5):230–241, 2002.

Motoo Kimura. Diffusion models in population genetics. Journal of Applied Probability, 1(2):177–232, 1964.

Motoo Kimura, Takeo Maruyama, and James F Crow. The mutation load in small populations. Genetics, 48(10):1303, 1963.

E Koch, NJ Connally, N Baya, MP Reeve, M Daly, B Neale, ES Lander, A Bloemendal, and S Sunyaev. Genetic association data are broadly consistent with stabilizing selection shaping human common diseases and traits. bioRxiv, 2024. doi: 10.1101/2024.06.19.599789.

Christopher C Kyriazis, Robert K Wayne, and Kirk E Lohmueller. Strongly deleterious mutations are a primary determinant of extinction risk due to inbreeding depression. Evolution letters, 5(1):33–47, 2021.

Russell Lande. The maintenance of genetic variability by mutation in a polygenic character with linked loci. Genetics Research, 26(3):221–235, 1975.

Russell Lande. Natural selection and random genetic drift in phenotypic evolution. Evolution, 30(2):314–334, 1976.

Russell Lande. Risk of population extinction from fixation of new deleterious mutations. Evolution, 48(5): 1460–1469, 1994.

Russell Lande and Susan Shannon. The role of genetic variation in adaptation and population persistence in a changing environment. Evolution, pages 434–437, 1996.

BDH Latter. Natural selection for an intermediate optimum. Australian Journal of Biological Sciences, 13 (1):30–35, 1960.

Michael Lynch, John Conery, and Reinhard Burger. Mutation accumulation and the extinction of small populations. The American Naturalist, 146(4):489–518, 1995a.

Michael Lynch, John Conery, and Reinhard Bürger. Mutational meltdowns in sexual populations. Evolution, 49(6):1067–1080, 1995b.

Susanne Paland and Bernhard Schmid. Population size and the nature of genetic load in gentianella germanica. Evolution, 57(10):2242–2251, 2003.

Roshni A Patel, Clemens L Weiß, Huisheng Zhu, Hakhamanesh Mostafavi, Yuval B Simons, Jeffrey P Spence, and Jonathan K Pritchard. Characterizing selection on complex traits through conditional frequency spectra. Genetics, 229(4):iyae210, 2025.

Nina Pekkala, K Emily Knott, Janne S Kotiaho, Kari Nissinen, and Mikael Puurtinen. The effect of inbreeding rate on fitness, inbreeding depression and heterosis over a range of inbreeding coefficients. Evolutionary Applications, 7(9):1107–1119, 2014.

Alan Robertson. The effect of selection against extreme deviants based on deviation or on homozygosis. Journal of Genetics, 54(2):236–248, 1956.

Jacqueline Robinson, Christopher C Kyriazis, Stella C Yuan, and Kirk E Lohmueller. Deleterious variation in natural populations and implications for conservation genetics. Annual review of animal biosciences, 11(1):93–114, 2023.

Jaleal S Sanjak, Julia Sidorenko, Matthew R Robinson, Kevin R Thornton, and Peter M Visscher. Evidence of directional and stabilizing selection in contemporary humans. Proceedings of the National Academy of Sciences, 115(1):151–156, 2018.

Guy Sella and Nicholas H Barton. Thinking about the evolution of complex traits in the era of genome-wide association studies. Annual review of genomics and human genetics, 20(1):461–493, 2019.

Yuval B Simons, Kevin Bullaughey, Richard R Hudson, and Guy Sella. A population genetic interpretation of gwas findings for human quantitative traits. PLoS biology, 16(3):e2002985, 2018.

Yuval B Simons, Hakhamanesh Mostafavi, Huisheng Zhu, Courtney J Smith, Jonathan K Pritchard, and Guy Sella. Simple scaling laws control the genetic architectures of human complex traits. PLoS biology, 23(10):e3003402, 2025.

Carl Veller and Pavitra Muralidhar. Quantitative system drift. Proceedings of the National Academy of Sciences, 123(12):e2527137123, 2026.

Bruce Walsh and Michael Lynch. Evolution and selection of quantitative traits. Oxford University Press, 2018.

Yvonne Willi, P Griffin, and J Van Buskirk. Drift load in populations of small size and low density. Heredity, 110(3):296–302, 2013.

Sewall Wright. The differential equation of the distribution of gene frequencies. Proceedings of the National Academy of Sciences, 31(12):382–389, 1945.

