## Supporting Information for "Mean fitness is maximized in small populations under stabilizing selection on highly polygenic traits"

Aaron P. Ragsdale

Department of Biology, University of Wisconsin–Madison, Madison, WI 53706

September 9, 2026

### Supplemental Information

#### A Accounting for environmental variance

We note that the following is standard, but include it here for completeness. We model phenotypes as $P = G + E$, where $E \sim \mathcal{N}(0, V_E)$ and is independent of $G$. Then $V_P = V_G + V_E$, and $V_E$ can be absorbed into $V_S$ in the fitness function by integrating over environmental backgrounds (Lande, 1975, and see Eq. A1 from Simons et al. (2018)):

$$w_e(z) = \frac{1}{(1 + V_E/V_S)^{1/2}} \exp\left(\frac{-(z-z_0)^2}{2(V_S+V_E)}\right). \tag{A1}$$

Setting $\tilde{V}_S = V_S + V_E$, this can be written as

$$w_e(z) = \left(\frac{V_S}{\tilde{V}_S}\right)^{1/2} \exp\left(\frac{-(z-z_0)^2}{2\tilde{V}_S}\right). \tag{A2}$$

Now the distribution of mean genetic values ($\bar{z}$) follows $\delta(\bar{z}) = \mathcal{N}\left(0, \frac{\tilde{V}_S}{2N_e}\right)$, and the distribution of genetic values follows $f(z|\bar{z}) = \mathcal{N}(\bar{z}, V_G)$. Then integrating over genetic values and the distribution of mean values (as in Eq. 3), we get

$$\begin{aligned}\mathbb{E}[\bar{w}] &= \int_{-\infty}^{\infty} \delta(\bar{z}) \int_{-\infty}^{\infty} w_e(z) f(z|\bar{z})\, dz\, d\bar{z} \\ &= \left(\frac{V_S}{\tilde{V}_S}\right)^{1/2} \left(\frac{2N_e\tilde{V}_S}{\tilde{V}_S + 2N_e(V_G + \tilde{V}_S)}\right)^{1/2}.\end{aligned} \tag{A3}$$

This takes the same form as Eq. 3 with $V_S$ replaced by $\tilde{V}_S$ and decreased by a factor

$$\left(\frac{V_S}{\tilde{V}_S}\right)^{1/2} = \left(\frac{V_S}{V_S + V_E}\right)^{1/2}.$$

Then, under the SHOC model,

$$V_G \approx \frac{4U\tilde{V}_S}{1 + \frac{\tilde{V}_S}{N_e\sigma^2}}.$$

Replacing this expression for $V_G$ above and carrying through the steps in the main text, the value of $N_e$ that maximizes $\mathbb{E}[\bar{w}]$ is unchanged (Eq. 8). Our condition for a fitness-maximizing finite $N_e$ is now

$$U > \frac{\sigma^2}{8\tilde{V}_S} = \frac{\sigma^2}{8(V_S + V_E)}. \tag{A4}$$

### B Predictions using the distribution of allele frequencies

When $V_G$ is no longer small compared to $V_S$, and for non-normally distributed effect sizes, the stochastic house-of-cards approximation (Eq. 5) may be a poor prediction for $V_G$. Instead, we may compute the expected additive genetic variance directly from the expected distribution of allele frequencies at trait-contributing loci. We assume a target size of $L$ loci and per-base mutation rate $\mu$, and that allelic effects for new mutations are drawn from some distribution $g(\alpha)$.

Assuming linkage equilibrium and additive effects, an allele with effect size $\alpha$ will be subject to underdominant-like selection, with selection coefficient $\frac{\alpha^2}{2V_S}(2x-1)$, with $x$ the allele frequency. Then the expected site-frequency spectrum, $\xi(x; N_e, \alpha)$, can be found using a variety of analytical or numerical methods, under either a recurrent mutation model (e.g., Wright, 1945; Barton, 1989), or an infinite-sites model (e.g., Kimura, 1964; Ewens, 2004, chapter 5). Here, we use a numerical implementation under the infinite sites assumption implemented in Jouganous et al. (2017).

The expected additive genetic variance is then calculated by integrating contributions from each effect size over over the distribution of effect sizes:

$$4N_e\mu L\int_{-\infty}^{\infty} g(\alpha)\alpha^2 \int_0^1 2x(1-x)\xi(x; N_e, \alpha)dxda.$$

When $V_G$ is non-negligible compared to $V_S$ ($V_G \sim 0.1 \times V_S$), the strength of selection for a trait-contributing allele is reduced, so that the selection coefficient is $\frac{\alpha^2}{2(V_S+V_G)}(2x-1)$. Computing the expected genetic variance using the SFS now depends on the genetic variance. In this case, we may iteratively compute distributions of allele frequencies to find expected $V_G$, and repeat this process until convergence, which is typically quick.

### C Simulations assuming linkage equilibrium

We consider a model where the dynamics of a trait-affecting allele depends on the mean genetic value and genetic variance in each generation. We assume linkage equilibrium is maintained at all times, avoiding the need to track individuals and the genotypes they carry. Thus, we only track the frequencies of segregating alleles, along with fixed alleles, and their effects. We assume new mutations occur at new sites and disallow back mutations.

Each generation, we compute the expected frequency of each allele in the next generation due to selection (described below), then apply binomial sampling to account for drift with the given population size. A Poisson-drawn number of new mutations are introduced (with mean $N_eU$), and their effects are drawn from a distribution $g(\alpha)$ symmetric around 0 with variance $\sigma^2$. In most cases we assume a normal distribution of effect sizes, though also explore a constant distribution (with all mutations having effect $\pm\alpha$ with equal probability) or an exponential distribution, again with positive or negative effects with equal probability (Figure S1). If a mutation reaches fixation at any point in the simulation, we continue to track its effect on trait values in the populations; lost mutations are discarded.

To compute the expected change in frequency for a given segregating allele (index $j$, with derived allele $A_j$ currently found at frequency $p_j$ and ancestral alleles $a_j$ with frequency $1-p_j$), we compute the expected difference in marginal fitness values $(w_{A_j} - w_{a_j})$ and use the formula $\Delta p_j = p_j(1-p_j)(w_{A_j} - w_{a_j})/\bar{w}$. We assume phenotypic values are normally distributed in the population with the mean and variance calculated over all variants ($\sum_i 2p_i\alpha_i$ and $\sum_i 2p_i(1-p_i)\alpha_i^2$, respectively). The mean fitness $\bar{w}$ is found by integrating against the Gaussian fitness function with the given optimum value and strength of selection $V_S$.

The marginal fitnesses are given by $w_{A_j} = p_j w_{A_jA_j} + (1-p_j)w_{A_ja_j}$ and $w_{a_j} = p_j w_{A_ja_j} + (1-p_j)w_{a_ja_j}$, where $w_{\cdot\cdot}$ is the average fitness of an individual carrying a given diploid genotype. To find $w_{A_jA_j}$, $w_{A_ja_j}$ and $w_{a_ja_j}$, we first calculate the background contributions to the mean and variance of genetic values from all other segregating and fixed alleles. These are $\bar{G}' = \sum_{i\neq j} 2p_i\alpha_i$ and $V_G' = \sum_{i\neq j} 2p_i(1-p_i)\alpha_i^2$. Then assuming a normal distribution of background trait values, the marginal fitness depends on the number of derived alleles $A_j$ carried by a given genotype. So for a genotype with $g_j \in \{0,1,2\}$ derived alleles, the marginal fitness is found by integrating $\mathcal{N}(\bar{G}' + g_j\alpha, V_G')$ against the Gaussian fitness function.

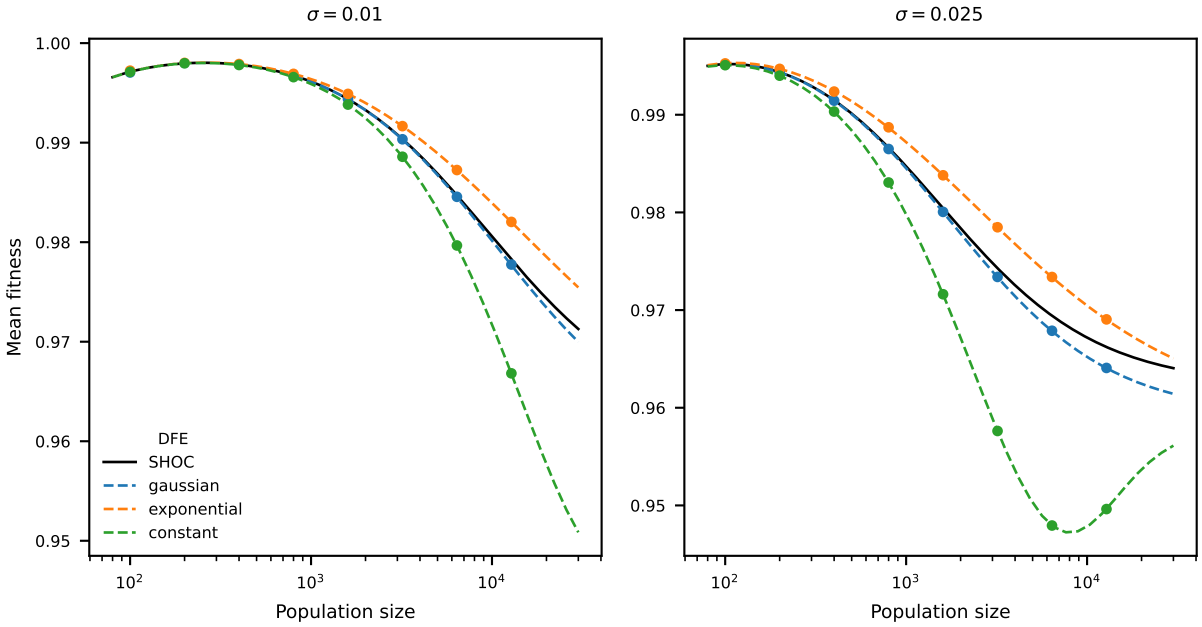


Figure S1: **Accuracy of the stochastic house-of-cards model for different distributions of effect sizes.** Using simulations assuming linkage equilibrium between all loci (Section C), we find that $V_G$, and thus $\bar{w}$, depends on the form of the distributions of effect sizes. Here, each distribution is symmetric with equivalent variances. The SHOC approximation is most accurate for normally distributed mutational effects, overestimates $V_G$ for an exponential distribution, and underestimates it for a "constant" DFE in which mutation effects $\alpha = \pm SD$ with equal probability. For the constant DFE, $V_G$ reaches a maximum (and $\bar{w}$ a minimum) around $4N_e\alpha^2 \approx 10V_S$, replicating the observation that the genetic variance is maximized for intermediate strengths of selection (e.g., see Figure 2 in Simons et al., 2018). Dashed lines represent predictions using the site frequency spectrum (Section B), which are highly accurate but are found numerically. In these simulations, the total diploid mutation rate $U = 0.02$ and the variance of new mutation effects from each distribution is made to be equivalent across the three distributions of effect sizes.

In all simulations assuming linkage equilibrium, we repeated simulations for each set of parameters 100 times. Each simulation replicate allowed a large number of generations for burn-in to explore the full stationary distribution of mean phenotypes. This requires more generations for small populations, and we used $10^4N_e$ for $N_e$ =100-200, $10^3N_e$ for $N_e$ =400, and $10^2N_e$ for $N_e \geq$800. After burn-in, we ran each replicate for $100N_e$ further generations, sampling population statistics every $N_e/10$ generations.

### D Individual-based simulations

Using the forward-in-time simulation software `fwdpy11` (Thornton, 2019), we simulated populations of varying size with different mutation and selection parameters. Parameter values were chosen to match those simulated assuming linkage equilibrium (Section C). Trait-affecting mutations could occur at any location in the chromosome (of length 1 Morgan) under an infinite-sites mutation model, and recombination rates were uniform across the entire region. Individual phenotypic values were found by summing across trait-affecting mutations they carried, and we assumed no environmental noise.

After allowing the simulation to run for at least $20N$ generations to reach quasi-steady state (and for many orders of magnitude larger for very small population sizes, see previous section), we stored the distribution of phenotypes and fitnesses across individuals every generation for $10N$ generations. From this information, we computed the average variances of phenotypes and mean fitnesses across all replicates of the same parameter set to compare to model predictions (Fig. 1). This was repeated 100 times, giving us 100 independent

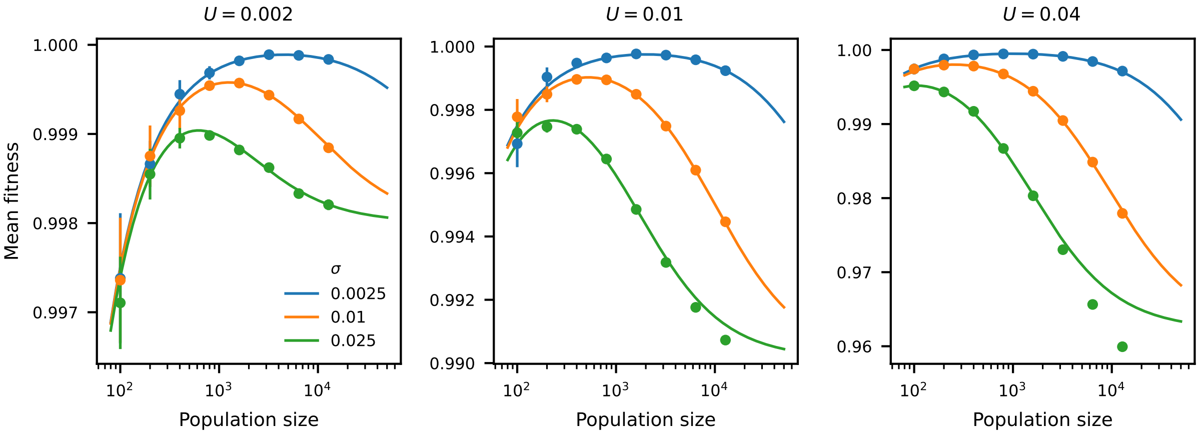


Figure S2: **Model predictions match individual-based simulations.** Repeating the set of parameters used to validate model predictions in Figure 1, we find good agreement between the SHOC-based predictions and simulations that account for linkage between loci. Error bars indicating 95% confidence intervals over replicates. $V_S$ is set to 1 in all simulations.

replicates. Within a given simulation, observed values from one generation to the next will be correlated, so these would not represent independent observations. To calibrate confidence intervals of measured quantities given the autocorrelation of measurements within replicates, we generated a set of bootstrapped values by resampling among replicates with replacement, taking the empirical inner 95% interval.

As we found with the simulations assuming linkage equilibrium, the individual-based simulations deviated from model predictions in parameter space where $V_G$ becomes non-negligible compared to $V_S$ (for larger population sizes and mutation rates). In this region of parameter space, polygenicity also increases, so that trait-affecting alleles are more densely distributed along the chromosome, leading to selective interference effects. The reduction of the effective strength of selection increases the genetic variance, and thus decreases mean fitness (beyond even what is observed in the simulations assuming linkage equilibrium). In other regions of parameter space, simulations again match predictions very closely. They also closely match the results from the simulations with LE, validating that simulation approach which is far more efficient than running individual-based simulations.

### E Pleiotropy

For multiple traits and pleiotropic selection, we may predict $V_G$ using the SFS under an appropriate scaling of the strength of selection. We assume an isotropic fitness function, noting that a change of variables can be found to transform a nonisotropic multivariate fitness function into one with equal strengths of selection and no covariance between transformed traits. However, under this transformation, mutational effects cannot be assumed to be isotropic. Nonetheless, the contributions of alleles with given multivariate effects can be computed, and then we may integrate over an arbitrary distribution of effects for new mutations.

$V_S$ is assumed to be equal for each trait (which we set to 1), with no covariance in the fitness function (i.e., an isotropic fitness model). The fitness function for multiple traits then becomes

$$w(\vec{z}) = \exp\left(-\frac{\|\vec{z}\|^2}{2V_S}\right).$$

We assume phenotypes are multivariate normally distributed, with means $\bar{z}_i$, variances $V_{G,i}$, and covariances $\text{Cov}(G)_{i,j}$. We approximate the population average phenotypes as having mean zero and variance $\frac{V_S}{2N_e}$. Then integrating the distribution of phenotypic values against the fitness function gives us the mean fitness.

Here, we consider a two-trait model. If trait values are uncorrelated, then

$$\mathbb{E}[\bar{w}] = \left(\frac{2N_eV_S}{V_S + 2N_e(V_{G,1} + V_S)}\right)^{1/2} \left(\frac{2N_eV_S}{V_S + 2N_e(V_{G,2} + V_S)}\right)^{1/2}$$

If the average phenotypic values are at the optima (i.e., $N_e \to \infty$) and phenotypes are possibly correlated, then

$$\mathbb{E}[\bar{w}] \approx \left(\frac{V_S^2}{(V_S + V_{G,1})(V_S + V_{G,2}) - \mathrm{Cov}(G_1, G_2)}\right)^{1/2}.$$

In our simulations, mutations are drawn from a multivariate normal distribution with zero means and arbitrary covariance matrix. We simulate two scenarios: one with no covariance between mutation effects, and another with nonzero covariance. In the simulations and model comparisons, the standard deviations of mutations for the two traits may be the same ($\sigma = 0.01$), or differ ($\sigma_1 = 0.005$ and $\sigma_2 = 0.02$). The correlation between effects on the two traits is set to either 0 or 0.8.

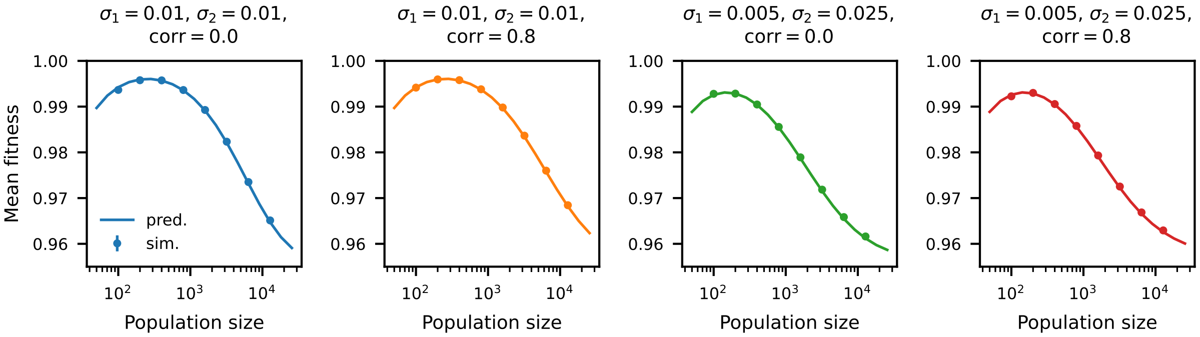


Figure S3: **Predictions match simulations for two traits and pleiotropic selection.** For two polygenic traits, either with the same or different variances of their distributions of effect sizes and with or without covariance in effects of an allele on the two traits, numerical model predictions using the SFS closely match simulations. In each case, the condition in Equation 9 is met for both traits. A more detailed study would examine arbitrary numbers of traits and scenarios of varying polygenicity and pleiotropy, but is beyond the scope of this work.

### F Adaptation following an optimum shift

Using the simulation framework assuming linkage equilibrium above (Section C), we simulate an instantaneous change in the optimum and track the evolution of the mean phenotype in the population towards that new optimum value. Here, the optimum shifts from $z_0 = 0$ to 1 (equal to one unit of $V_S$, which is set to 1 in all simulations). Small populations harbor lower genetic variance, and therefore adapt more slowly to the new optimum (Lande, 1976), and this is observed in simulations (Figure S4).

### References


Nick Barton. The divergence of a polygenic system subject to stabilizing selection, mutation and drift. *Genetics Research*, 54(1):59–78, 1989.

Warren John Ewens. *Mathematical population genetics: theoretical introduction*, volume 27. Springer, 2004.

Julien Jouganous, Will Long, Aaron P Ragsdale, and Simon Gravel. Inferring the joint demographic history of multiple populations: beyond the diffusion approximation. *Genetics*, 206(3):1549–1567, 2017.

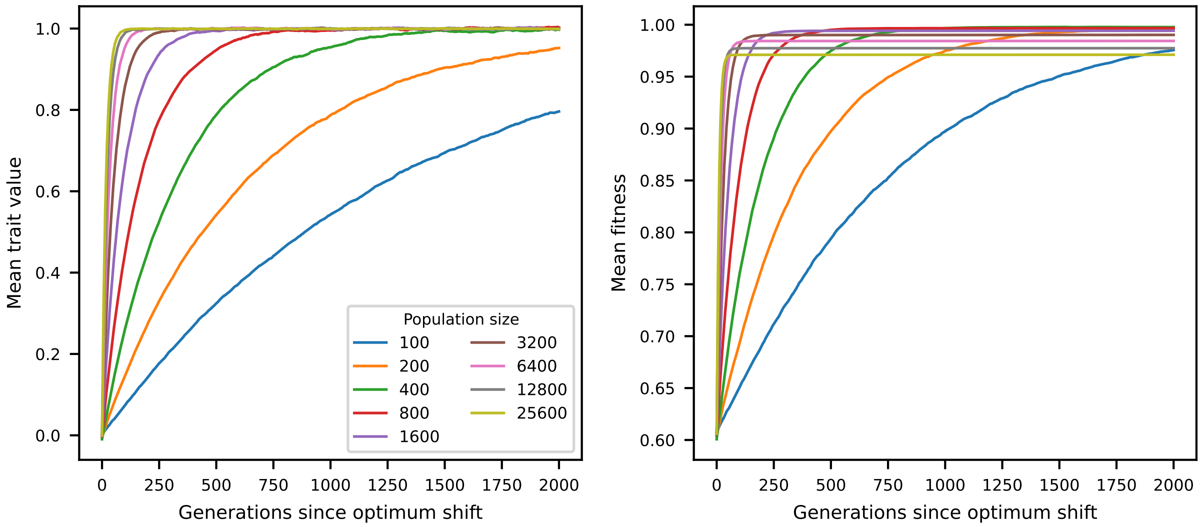


Figure S4: **Small populations more slowly approach a new optimum.** Due to lower genetic variance contributing to the trait, the rate of adaptation to a new phenotypic optimum is reduced in small populations compared to large populations. In these simulations, assuming perfect linkage equilibrium between trait-contributing alleles (Section C), $U = 0.04$ (total diploid rate), $\sigma = 0.01$, $V_S = 1$ and the optimum shifts instantaneous from $z_0 = 0$ to 1. The mean trait value and fitness were tracked for 2000 generations post-optimum shift, and simulations were repeated 100 times for each set of parameters.

Motoo Kimura. Diffusion models in population genetics. *Journal of Applied Probability*, 1(2):177–232, 1964.

Russell Lande. The maintenance of genetic variability by mutation in a polygenic character with linked loci. *Genetics Research*, 26(3):221–235, 1975.

Russell Lande. Natural selection and random genetic drift in phenotypic evolution. *Evolution*, 30(2):314–334, 1976.

Yuval B Simons, Kevin Bullaughey, Richard R Hudson, and Guy Sella. A population genetic interpretation of gwas findings for human quantitative traits. *PLoS biology*, 16(3):e2002985, 2018.

Kevin R Thornton. Polygenic adaptation to an environmental shift: temporal dynamics of variation under gaussian stabilizing selection and additive effects on a single trait. *Genetics*, 213(4):1513–1530, 2019.

Sewall Wright. The differential equation of the distribution of gene frequencies. *Proceedings of the National Academy of Sciences*, 31(12):382–389, 1945.
